# Evaluation of antimicrobial activity of Kefir drink against Escherichia coli, Salmonella typhimurium and Shigella flexneri

**DOI:** 10.1101/2023.02.22.529598

**Authors:** Zaira Cecilia Gutiérrez Cortéz, C. Alonso Rubén Tescucano Alonso, Jorge Angel Almeida Villegas, Gabriel Martínez González

## Abstract

**Objective:** Evaluate the antimicrobial activity of the Kefir drink against *Escherichia coli, Salmonella typhimurium and Shigella flexneri*

**Method:** The Kefir used in the investigation was acquired in Toluca, State of Mexico. We performed a reactivation of Kefir with pasteurized milk and analyzed 3 carbohydrates (honey, sugar and piloncillo) at different concentrations and times, 80, 100, 120% and 24, 48 and 72 hours respectively. Isolated cultivable microorganisms were characterized by phenotypic, biochemical and mass spectroscopy techniques. The initial and final pH were determined during the study time. The antimicrobial activity was carried out by extracting the metabolites present in the ferment with the Kirby-Bauer method, in addition the direct ferment was evaluated, to determine if there was inhibition with the *Escherichia coli (*ATCC 11229) strains, *Salmonella typhimurium* (ATCC 14028) and *Shigella flexneri* (ATCC 12022).

**Results:** It was observed that in the three carbohydrates used at a concentration of 120% and at a time of 72 h, a lower pH was obtained (3.51 to 3.64) compared to their initial concentration (6.50 to 6.64). From the metabolites extracted in the different ferments, no inhibition halo was obtained with the strains analyzed. However, when using direct ferments, it was observed that in the carbohydrates used (sugar, honey, piloncillo) there was the presence of an inhibiting halo or the growth of colonies other than those evaluated. The isolated cultivable microorganisms were: *Pichia kudriavzevii* (yeast); *Enterococcus* sp (gram-positive coconut) and *Lactobacillus* sp (gram-positive bacillus).

**Conclusions:** Kefir ferments made with different carbohydrates, came to present a degree of inhibition only as a consortium against Gram-negative microorganisms analyzed.

## Introduction

Gastrointestinal diseases are one of the main public health problems, these diseases are caused by bacteria (mainly *Escherichia coli, Salmonella* and *Shigella*), parasites and viruses. Likewise, these types of infections are among the most frequent infectious diseases. (1)

The trend in Mexico on salmonellosis in the period from 2000 to 2012 is upward with respect to notifications, of 107,289 and 183,043 cases, respectively. There is a maximum peak in 2019 with 185,290 cases and as of 2013 a downward trend is observed. On the other hand, it is estimated that *Shigella* causes approximately 600,000 deaths per year worldwide since its distribution is worldwide. In the year 2021, 1,992 cases were reported. Lastly, enteropathogenic *Escherichia coli* frequently causes epidemic outbreaks and has been associated with high mortality (10-40%). In 2010, 121,455 deaths were reported from this pathotype. (2)(3)

Kefir is a fermented milk product derived from the Turkish word keyif, which means “feel good”. Other names by which kefir is known are: kefyr, kefir, kefer, kiaphur, knapon, kepi or kippi. The origins of kefir are found in the mountains of the Caucasus, where in ancient times it was characterized by the breeding of goats that provided milk for consumption through milking, ensuring food when production was scarce. Kefir granules are made in the same region and are still made completely by hand. It has been widely used in Russia and Central Asian countries such as Kazakhstan and Kyrgyzstan for centuries, and is now gaining popularity in European countries, Japan and the United States for its nutritional and therapeutic benefits. (4)(5)

Kefir is fermented milk produced from grains that comprise a specific and complex mixture of bacteria and yeasts that live in a symbiotic association (6). The grains have a jelly and translucent appearance, yellowish to brown, with irregular shapes and sizes ranging from millimetres to a few centimetres.(7) kefir has been associated with a range of health benefits such as cholesterol metabolism and angiotensin-converting enzyme (ACE) inhibition, antimicrobial activity (8). Probiotics stand out among fermented foods. According to the definition, probiotics are living microorganisms that provide health benefits to those who consume them.(9)

The grains contain lactic acid bacteria (LAB) such as lactobacilli (in a composition of 65-80% depending on their preparation), lactococci, leuconostoc, streptococci, as well as some yeasts (*Candida* sp., *Kluyveromyces* sp., *Saccharomyces* sp., *Torulopsis* sp., *Zygosaccharomyces* sp.), sometimes acetic acid bacteria (*Acetobacter* sp.), casein, and complex sugars.(10) *Bifidobacterium* sp., *Lactobacillus* sp. and probiotic yeast (*Saccharomyces boulardi*i) may be used as adjunct cultures when blended with kefir grains or kéfir cultures.(11) Traditional kefir contains *Saccharomyces cerevisiae, Pichia fermentans, Kazachastania unispora*, and *Kluyveromyces marxianus* and lactic in addition to many other smaller populations of yeast, the commercial variant only has *Saccharomyces cerevisiae*. (12)

## Material and methods

### Activation of Kefir grains

The grains of Kefir were acquired in Toluca, Mexico and for its activation was performed according to the modified methodology of Macuamule et al. (13), where commercial milk was pasteurized at 80°C for 15 min, with constant agitation and allowed to cool at room temperature, Once this was done, the kernels of Kefir were placed and incubated at 25 °C for 24 h, leaving the lid unsealed, as there is presence of gas. After the incubation time, the grains were sieved and placed again in double pasteurized milk, this process was repeated for 4 consecutive days. (13)

After the days of activation of Kefir grains, from two base formulations according to Macuamule et al. (13) and Monar et al. (14), a formula was made for the preparation of the samples to be studied, which was established as 100% having the following formulation: 95 mL of pasteurized milk; 4.27 g of carbohydrate to analyze (sugar, honey or piloncillo) and 1.71g of the active grain of Kefir. Subsequently, calculations were made to obtain concentrations of 80% and 120%. Sugars were acquired in the municipality of Toluca, Mexico. Three fermentation times (24h, 48h and 72h) were considered, for each formulation made (80, 100 and 120%), obtaining 27 combinations with the three carbohydrates to analyze. Once the samples were prepared, the pH was measured as the starting point and incubated at 35 °C± 2°C for the established fermentation times.

### Extraction of metabolites obtained by fermentation

After the fermentation time, two samples were taken; one to isolate the microorganisms present and the other to extract the metabolites present in the fermentation carried out.

To extract the metabolites present in the fermented samples, an aliquot was centrifuged at 14 000 rpm for 10 min and the supernatant was transferred to a new Falcon tube with the aid of a sterile syringe, then centrifuged again under the same conditions. The supernatant was collected and filtered with 0.22 µm syringe filters (GVS), deposited in sterile test tubes at 4°C.(13) Microorganisms used.

The following microorganisms were used: *Escherichia coli (*ATCC 11229), *Salmonella typhimurium* (ATCC 14028) and *Shigella flexneri* (ATCC 12022), belonging to the biotery of the University of Ixtlahuaca-CUI. The microorganisms were lyophilized and activated when reseeded in trypticase soy agar and incubated at 35°C+-2°C per 24h. Over time, its growth and purity were verified through a Gram stain. After confirming that they were Gram-negative bacilli, they were reseeded in various biochemical tubes (LIA, TSI, MIO, malonate broth, Simmons citrate, urea broth), to corroborate genus and species of each microorganism.

### Evaluation of antimicrobial activity of metabolites extracted in fermentation (Kirby-Bauer)

Each test microorganism, it was grown by cross stretch marks, in Trypticase Soy agar (TSA) (DIBICO), incubating at 37 °C for 24 hours, from the individual planting colonies were taken, placed in a tube with sterile saline, matching turbidity with standard 0.5 Mc Farland (1.5×10^8^ UFC/mL) tube. From each standardized inoculum, it was sown by closed stria with the help of a sterile swab in a box of Müeller Hinton agar (DIBICO), then placed the discs grade AA with sterile tweezers distributed on each petri box, and placing 10 μL of the different metabolites extracted from the 27 combinations obtained. (15)

The antibiogram of each microorganism used in the research was also performed, with the following antibiotics: Ampicillin 10 μg (BD BBL); Chloramphenicol 30 μg (BD BBL); Ciprofloxacin 5 μg (BD BBL); Amoxicillin-clavulanic acid 20/10 μg (BD BBL) and Polymyxin 300 UI (BBL).

Each trial was performed in triplicate. The results of antibacterial activity were analyzed based on the statistical program SPSS, by means of an ANOVA test (Analysis of variance), comparison of Tukey means (p 0.05).

### Evaluation of the antimicrobial activity of the Kefir ferment obtained

To evaluate the antimicrobial activity of Kefir ferment with the different carbohydrates evaluated, a suspension adjusted to 0.5 Mac Farland (1.5×10^8^ CFU/mL) was used individually, considering the strains with a reseeding of 20h and a massive seeding was performed under aseptic conditions in Tryptecasein Soy agar (TSA) with the support of a sterile swab.

In each TSA box with the inoculated microorganism, three single streaks were deposited vertically, using the 80%, 100% and 120% ferments with each carbohydrate used at 48 and 72h times. The plates were incubated inverted at 35°C ± 2°C and visualized every 24h for 3 days.

### Isolation of the cultivable microorganisms present in the Kefir ferment in LBS agar

From the microbial growth observed in the single stria space of the results of the evaluation of the antimicrobial activity of the Kefir ferment obtained, a roast was taken and the colonies observed in LBS agar (Lactobacillus Selection Agar) a 35°C 2°C per 24 h. After incubation time, 21 zones with inhibition halo were selected to perform a Gram stain and select strains with different morphology for identification and isolation. Gram staining was also performed directly from the ferments with the different carbohydrates evaluated.

## Results

The initial pH of the different preparations ranged from 6.50 to 6.64. The results of the ferments with the different carbohydrates were as follows: 1) regarding sugar, the lower pH was 3.64 at a concentration of 120% and 72 h; 2) according to honey, the lower pH was 3.61 at a concentration of 120% and 72 h; 3) regarding the piloncillo, the lowest pH was 3.52 at a concentration of 120% and 72 h (see Graph 1). It was observed that in the three carbohydrates used at a concentration of 120% and in a time of 72 h, was the situation where a lower pH was obtained, with a decrease of 45 %, compared to the initial concentration.

### Evaluation of antimicrobial activity of metabolites extracted in fermentation (Kirby-Bauer)

From the metabolites extracted in the different ferments, no inhibition halo was obtained with the strains analyzed. However, inhibition halos could only be observed with the sensidisks used with antibiotics, which when compared with the CLSI tables could be observed the following: 1) With *Shigella flexneri* ATCC 12022, it was sensitive to ampicillin, chloramphenicol, ciprofloxacin, amoxicillin-clavulanic acid and resistant to polymyxin; 2) Regarding *Escherichia coli* ATCC 11229, it was sensitive to chloramphenicol, intermediate with: ampicillin, ciprofloxacin, amoxicillin-clavulanic acid and resistant to polymyxin; 3) With *Salmonella typhimurium* ATCC 14028, was sensitive to chloramphenicol, ciprofloxacin, amoxicillin-clavulanic acid and resistant to ampicillin and polymyxin.

### Evaluation of the antimicrobial activity of the Kefir ferment obtained

From the results of the evaluation of antimicrobial activity, it was observed that, in all ferments with sugar carbohydrates, honey, piloncillo and with the presence of microorganisms of *Escherichia coli* ATCC 11229, *Salmonella typhimurium* ATCC 14028 and *Shigella flexneri* ATCC 12022, there was the presence of a halo of inhibition or in the growth of colonies within the stria that was performed with the ferment, as seen in Figure 1.

**Graph No. 1.**
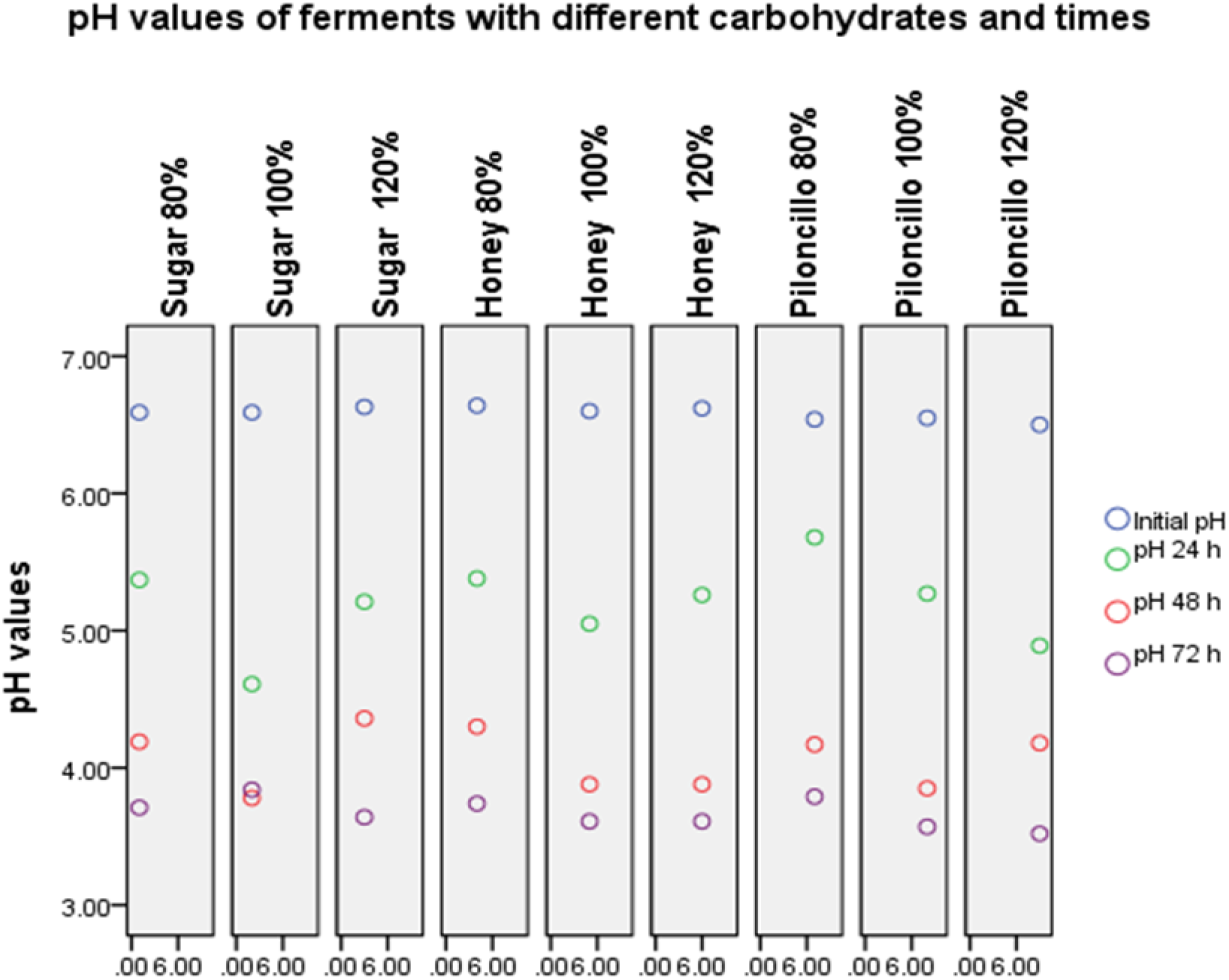
Determination of the pH of the different ferment preparations with sugar, honey and piloncillo at an initial pH, 24 h, 48 h and 72 h. Source: Own.

**Figure 1.**
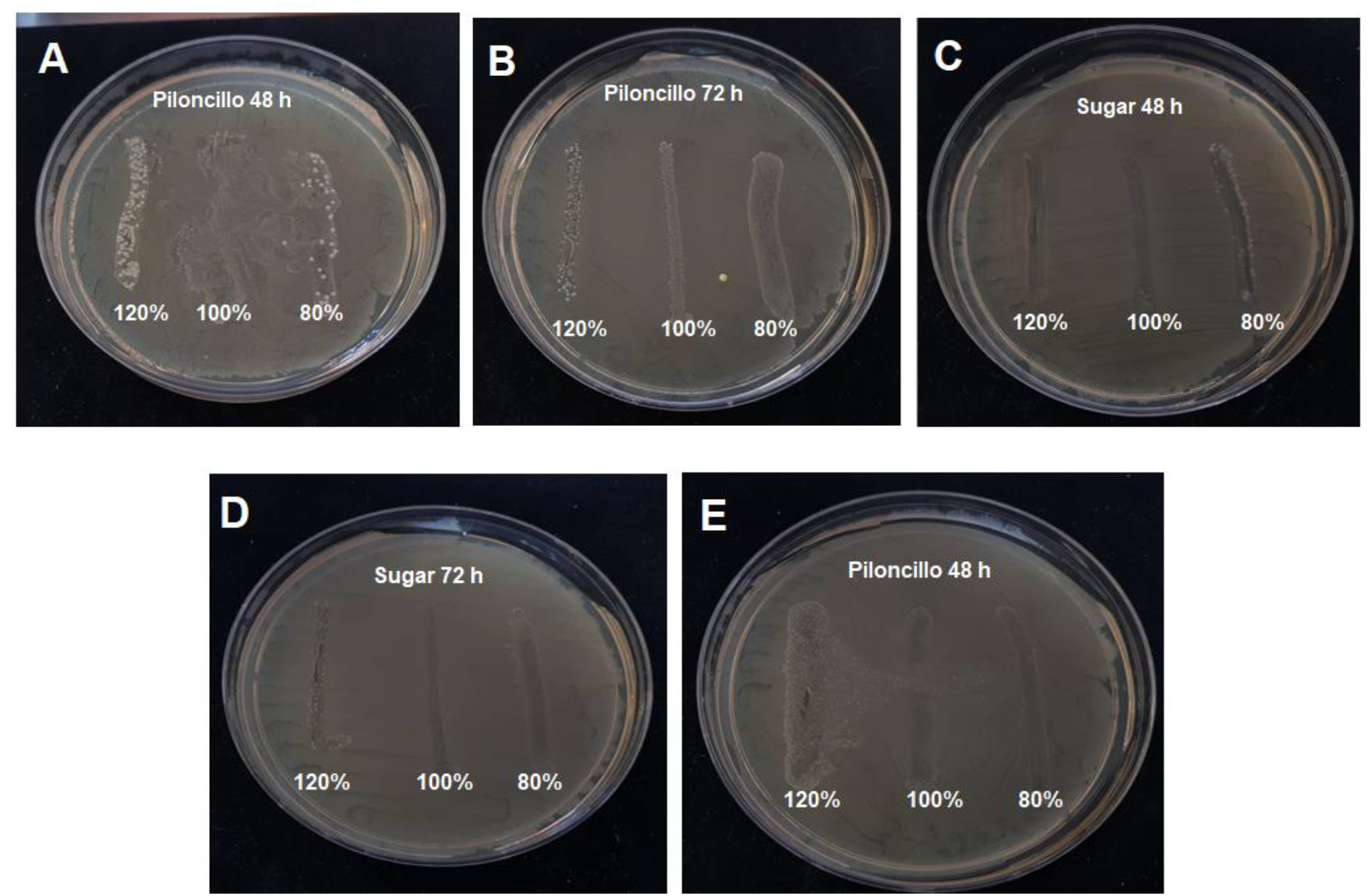
Results of the evaluation of the antimicrobial activity of Kefir ferment with piloncillo and sugar. In the A-E images, the presence of inhibiting halos is observed in the single stretch marks performed, in addition to the growth of colonies within the halo produced. Images A and B were evaluated against *Shigella flexneri* ATCC 12022 and images C to E with *Escherichia coli* ATCC 11229. Source: Own.

### Isolation of the cultivable microorganisms present in the Kefir ferment in LBS agar

From the various inhibition halos observed in the boxes with the microorganisms and the sugar, honey and piloncillo ferments, a Gram stain was made, to see which morphology of microorganisms was observed and then could be isolated and identified. It was possible to determine the presence of three microorganisms, which were: a Gram positive bacillus, a Gram positive cocci and a yeast. The only ferment, where there was the presence of two microorganisms (a cocci and a yeast), was with the Piloncillo at a concentration of 120%/72h and with *Shigella flexneri (*ATCC 12022). Of the other isolates in the various ferments, only one microorganism was present and the pathogen was no longer observed (*Escherichia coli* ATCC 11229, *Salmonella typhimurium* ATCC 14028 or *Shigella flexneri* ATCC 12022).

Gram staining was also performed taking an aliquot of the direct ferments of the three carbohydrates without adding the pathogens to be evaluated, observed in the same way the presence of the three different morphologies observed previously and with greater quantity the Gram bacillus (+).

The three isolated microorganisms were sent to the MICRO-TEC laboratory for identification, using automated equipment (MicroScan autoSCAN-4) by Beckman Coulter and spectrometry with a MALDI TOF, giving the following results: *Pichia kudriavzevii* (yeast); *Enterococcus sp* (Gram positive cocci) and *Lactobacillus* sp (Gram positive bacillus).

## Discussion of results

Fermented dairy products are ideal for delivering probiotic bacteria to the human intestine and providing the optimal environment for growth. Probiotics are described as “living microorganisms that provide a health benefit to the host when provided in sufficient doses” (16). Recently, consumer interest in fermented products containing probiotic microorganisms has increased, demonstrating the regulatory effects of kefir microorganisms on the gut microbiota and its accumulated antibacterial activity. (17)

Kefir is a fermented dairy product derived from the Turkish word keyif, which means “feel good”. Other names by which kefir is known are: kefyr, kefir, kefer, kiaphur, knapon, kepi or kippi. The origins of kefir are found in the Caucasus mountains. Previous studies on kefir have reported antibacterial, immunological, antitumor, cholesterol-lowering and β-galactosidase activity. (4)

The kefir used in the research was acquired in Toluca, State of Mexico and was activated with double pasteurized milk, obtaining hard grains, irregular in shape, and yellowish-white, having a cauliflower type form, being the traditional form found in grains, in addition other forms of lamina type can be found, nucleus type structures and robust structures with little undulation. In an investigation conducted by Kalamaki and Angelidis (18), from 12 samples of Kefir grains from Greece, they most frequently observed the type of cauliflower (8 of 12 samples).

In the present research work, the carbohydrates used decreased the initial pH by up to 45%, after 72h with a concentration of 120%, this due to the presence of the microorganisms that make up the Kefir consortium. Within the isolated microorganisms, there is the presence of *Lactobacillus* sp and *Enterococcus* sp, which are part of the lactic acid bacteria (LAB) and the yeast *P. kudriavzevii* (19). Kefir grains are composed of a complex symbiotic microbial ecosystem of lactic acid bacteria (LAB), acetic acid bacteria, and yeast, embedded in an exopolysaccharide matrix known as kefiran, which consists of approximately equal proportions of D-glucose and D-galactose, which protects the grain microbiota from adverse environmental conditions and represents between 24 and 25% of the dry weight of the grain of kefir. (20)

The pH reported by some authors for lactic acid bacteria ferments ranges between 3.8 and 4.5, which coincides with the values obtained with the microbial consortium. In our study, a value of 3.52 with greater acidity was obtained due to the effect of fermentation caused by *P. kudriavzevii*. Lactic acid bacteria are a kind of microorganisms that can ferment carbohydrates to produce lactic acid. Lactic acid bacteria can decompose macromolecular substances in food, including degradation of indigestible polysaccharides and transformation of undesirable flavor substances. This explains the decrease in pH due to the fermentation of the additional sugars used in milk cultures. And a pH lower than that reported in the literature. Bacteriocins and hydrogen peroxide produced by LAB could inhibit the growth of *E. coli* and other pathogens. Another mechanism of LAB is to enhance the systemic and mucosal immune response to improve the health of the body. (21)(22)

On the other hand, lactic acid bacteria have been found to have another health benefit, as they are related to improving the pathophysiology of metabolic, neoplastic and infectious diseases, which includes inflammation, all this by releasing molecules charged with the activity of immunomodulation. In this regard, it is reported that the acidity produced in fermentation or by the same microbiota, could be influencing the immune response in inflammation by interfering with the nuclear factor pathway Kappa B, by releasing inflammatory mediators, That is, the pathway can be regulated by the biological effect of the secreted molecules or by the change of pH, consequently, modulating the expression of this transcription factor, generates an anti-inflammatory environment that is necessary to avoid the diseases already mentioned (20).

On the other hand, *Enterococcus* are classified as homo and heterofermentative bacteria, producing up to 85% lactic acid and the rest ethanol and other acid derivatives. For *Lactobacillus* they are mainly heterofermentative and have a different mechanism to ferment carbohydrates, since they can be heterolactic or homolactic, depending on the species. In the case of *Lactobacillus*, the Embden Meyerhof glycolytic pathway usually predominates. Both bacteria isolated in the kefir under study are responsible for the decrease in pH. As shown in Graph No. 1 On the other hand *Pichia kudriavzevii* is a yeast with fermentative characteristics that uses glucose to produce ethanol and also the ability to produce lactic acid under stress situations such as low pH, in addition to other substrates to reach to the same compound.(23)(24)(25)

In the evaluation of the antimicrobial activity of the metabolites extracted in fermentation with the different carbohydrates analyzed, no inhibition halos were observed with the pathogenic microorganisms tested at a concentration of 1.5×10^8^ CFU/mL, so, it is suggested that it could be that the extracted metabolites were not in sufficient quantity to act against pathogens or as established by the work of González et al.(20) that considering that there are several kefir supernatants and these contain different inhibitory compounds, these compounds may interact with each other to improve or antagonize their antibacterial effects, being an example of this mechanism, inactivation of bacteriocins by organic acids or enzymatic degradation during fermentation.

However, when the antimicrobial activity was evaluated with the microorganisms present in the grain of kefir and with their metabolites present in the ferments, inhibition halos and growth of other colonies than the pathogenic microorganisms evaluated were observed. These different colonies when performing a Gram stain were: a bacillus Gram (+), a coconut Gram (+) and a yeast, also when sent to identify phenotypically these were: *Lactobacillus* sp, *Enterococcus* sp and *Pichia kudriavzevii*, respectively.

In reviewing the inhibition halos, only the presence of a single microorganism could be observed in most halos, for example with the Piloncillo ferment: 1) with a concentration of 120% and at 48h in the presence of *Salmonella typhimurium* (ATCC 14028), the presence of *Lactobacillus* sp; 2) with a concentration of 120% and at 48h in the presence of *Escherichia coli* ATCC 11229 could be observed the presence of *Lactobacillus* sp; 3) with a concentration of 120% and at 72h in the presence of *Shigella flexneri* ATCC 12022, the presence of *Pichia kudriavzevii* was observed; 4) with the concentration of 80% and 120% at 48h in the presence of *Shigella flexneri* ATCC 12022, the presence of *Enterococcus* sp.

The only ferment where the presence of *Pichia kudriavzevii* and *Enterococcus* sp was observed was with the Piloncillo at a concentration of 120%/72h and with *Shigella flexneri* ATCC 12022, establishing a synergistic effect. Sarkar (26) states that yeasts and lactobacilli are mutually dependent and grow in balanced proportions within kefir grains, and a symbiosis has been observed between yeasts, lactobacilli and streptococcus during kefir production.

The antibacterial activity of individual isolates of kefir has been extensively studied in vitro (20). Kakisu et al. (27) showed that *Lactobacillus plantarum* CIDCA 8311 exhibited potential inhibitory properties against species that cause Shigellosis disease (*Shigella flexneri* and *Shigella sonnei*) in Hep-2 cell lines. Santos et al. (28) evaluated the antibacterial activity of 58 strains of the family Lactobacillus isolated from kefir, and about 75% of the strains showed antibacterial activity against the enterohemorrhagic strain of *Escherichia coli* and *Yersinia enterocolitica*; among them the strains of *Lactobacillus acidophilus* CYC 10050 and CYC 10051 and *Lactobacillus kefiranofaciens* CYC 10058 inhibited *E. coli* CECT 4076, *Listeria monocytogenes, Salmonella typhimurium, Salmonella enteritidis, Shigella flexneri* and *Yersinia enterocolitica*.

The genus *Enterococcus* has been shown to produce bacteriocins, which have demonstrated antimicrobial activity against pathogenic microorganisms, such as: *E. coli* and *Salmonella* sp. On the other hand, *Pichia kudriavzevii* has been reported to produce inhibition halos of 8.56 and 9.50 mm for *E. coli* and *Salmonella* sp, demonstrating that it produces antimicrobial substances capable of counteracting these pathogens in vitro. (29)

The antibacterial activity of kefir can be due to a variety of factors, starting with proteolytic enzymes, organic acids, CO_2_, bacteriocins, bioactive peptides, as well as competition for nutrients and the generation of antibacterial compounds during carbohydrate metabolism by lactic acid bacteria. In addition, decreased pH of kefir due to accumulation of organic acids leads to broad-spectrum inhibitory activity against Gram-positive and Gram-negative bacteria. (20)

It has also been seen that the genera present in the Kefir grain microbiota, how those found in research, are similar to the microbiota in the human, therefore, research has shown that they can generate and secrete precursors of certain chemical messengers, such as tyrosine, which is important in the formation of catecholamines; or tryptophan, serotonin precursor, even release neurotransmitters that add to the functioning of the immune system. Undoubtedly, in this context, more studies are needed to know in depth the interaction of these microorganisms with the individual, and their modulating effect on the immune system or other systems. (30)

## Conclusions

It was observed that in the three carbohydrates used at a concentration of 120% and at a time of 72 h, it was the situation where a lower pH was obtained (3.51 to 3.64), than their initial concentration (6.50 to 6.64). From the metabolites extracted in the different ferments, no inhibition halo was obtained with the strains analyzed. However, when using direct ferments, it was observed that in the carbohydrates used (sugar, honey, piloncillo) there was the presence of an inhibiting halo or the growth of colonies other than those evaluated. The isolated cultivable microorganisms were: *Pichia kudriavzevii* (yeast); *Enterococcus* sp (gram-positive coconut and *Lactobacillus* sp (gram-positive bacillus). It is considered that more experiments should be done to determine which mechanisms are attributed to inhibition.

## Acknowledgment

The University of Ixtlahuaca CUI is thanked for the support provided and all the members of the Bachelor’s Degree in Biological Pharmaceutical Chemistry who helped in the realization of the work

## Conflict of interest

The authors declare that they have no conflict of interest.

## Notes

### Competing Interest Statement

The authors have declared no competing interest.

